# Injectable Immune-Engineered Hydrogel Niche Remote From The Immune Suppressed Tumor Microenvironment For Cancer Immunotherapy

**DOI:** 10.1101/2025.05.11.653292

**Authors:** Saira Nujoom Muhammad, Zahara T Zakariya, Sherin Shaji, Anjali K Sunilkumar, Amal George, Sreedevi P Radhakrishnan, Shantikumar V Nair, Manzoor Koyakutty

## Abstract

Immunocompromise is a hallmark of cancer, affecting both the peripheral immune system and local tumor microenvironment (TME). Current immunotherapies like checkpoint inhibitors, CAR-T cells, and neo-antigen vaccines show limited efficacy due to severe immunosuppression in most patients. Here, we report an immunologically engineered injectable nano-hydrogel (iHG) that can: (i) recruit the desired set of immune cells away from the suppressed TME and peripheral organs, (ii) activate them within a protective ambit of engineered immune-stimulatory hydrogel niche, and (iii) release them to target cancer even in distant locations. Biodegradable and injectable iHG compositions were tested and optimized for their ability to attract and activate dendritic cells, macrophages, monocytes, NK cells, B cells, and T cells via STING, TLR, CD86, and Th1-polarized cytokine pathway without requiring exogenously introduced neo-antigens as vaccines. In a mouse melanoma model, optimized iHGs elicited a robust antitumor immune response through innate and adaptive arms. Most importantly, iHGs as a single agent immunomodulator exhibited better tumor control than when combined with anti-PD1 immune checkpoint antibody. These findings highlight the potential of engineering immunologically functional and injectable hydrogel niches as a new type of immunotherapeutics to reprogram immune cells to overcome both local and systemic immunosuppression and combat cancer effectively.

## 1 INTRODUCTION

Cancer is a disease marked by oncogenic mutations and immunosuppression, which together drive its progression [1]. Cancer cells not only modify their primary site by creating a favourable stroma and neo-vascularization, but also systemically reprogram both circulating and tissue-resident immune cells to support their growth.[2] As tumors become clinically detectable, they adopt mechanisms akin to peripheral immune tolerance.[3,4] A hallmark of this systemic manipulation is the recruitment of immunosuppressive cells to the tumor microenvironment (TME).[2] Ultimately, the entire body of a cancer patient can transform into a tumor-supportive “macroenvironment,” where cancer cells evade immune surveillance, proliferate, circulate, and metastasize. This is brought about for example in several ways: formation of immunosuppressive premetastatic niches systemically; by circulating tumor cells (CTCs) that can stimulate neutrophils to exert immunosuppressive effects; or by secretion of extracellular vesicles from CTCs that can induce apoptosis in cytotoxic T cells, inhibit natural killer (NK) cells and promote the expansion of regulatory T cells (Tregs).[5–7] Paradoxically, cancer exploits immune hubs like lymph nodes and the bloodstream to spread to distant organs, underscoring the profound immunosuppression present throughout the systemic organs, in addition to local TME.[4,8]

The profound role of immunosuppression in cancer is evidenced by the limited success of immune checkpoint inhibitors (ICIs) and other immunotherapeutics.[9–11] Monoclonal antibodies targeting PD-L1/PD-1 or CTLA-4 have demonstrated significant efficacy in a subset of patients[12–16], while CAR-T therapies targeting CD19 in B-cell have shown great promise in leukaemia but largely failed against solid tumors.[17,18] Clinical data reveal that only < 25% of patients benefited from all forms of cancer-immunotherapeutics, and 75% showed only partial or no response, primarily due to the prevailing immunosuppression not only in the tumor microenvironment (TME) but the whole body.[9,19–25] It is now recognized that successful cancer immunotherapy requires neutralizing both systemic and local immunosuppression.[26–28]

To address systemic and local immunosuppression, here we propose a novel strategy: an exogenously administered, immune hydrogel (iHG) engineered niche designed to recruit and activate immune cells against cancer at a site distant from the immunosuppressive tumor microenvironment and peripheral organs. This engineered nanomaterial hydrogel niche provides an immune-activation platform free from pre-existing suppressive conditions, enabling immune cells to assemble, activate, and effectively target cancer. This injectable and biodegradable immune-niche can be repeatedly administered to periodically recruit and activate immune cells, as the initially recruited cells may get exhausted during the continuous engagement with cancer. Effectively, iHG will act as a temporary ‘immune-cell camp’ mimicking native lymphoid structures, enabling immune cell assembly, interaction, training, expansion, and dispersal into the systemic circulation to combat cancer. [29] This approach reprograms antitumor immunity in a manner akin to extra nodal tertiary lymphoid structures (TLSs)[30–33]

Biomaterial hydrogels were reported as localized platforms for creating artificial TLS, employing scaffolds to attract immature dendritic cells (DCs) toward cancer vaccination.[29,33–35] These scaffolds can locally present adjuvants and antigens, triggering DC activation, antigen processing, and presentation. [36–40] David Mooney et al. have developed various implantable biomaterial niches for *in situ* vaccination against cancer.[33,39] Similarly, injectable mesoporous silica rods have been explored as cellular microenvironments for *in situ* cancer vaccination.[41] Xiao-Kang Jin et al recently reported STING-activating metal ion complexed hydrogels that mimic tertiary lymphoid structures as anti-cancer immunotherapeutics.[42] Similarly, an injectable hydrogel system using L-norvaline-containing di-block copolymer that self-assembles into a thermo-responsive gel was reported by Xiaomeng et al. [43] This hydrogel effectively blocks the arginase-1 (ARG1) pathway and modulates the tumor microenvironment. Additionally, pore-forming hydrogel systems incorporating porogen-dispersed alginate networks have been shown to recruit dendritic cells and activate T cells in situ through the controlled release of GM-CSF and the IDO inhibitor thereby reshaping the immunosuppressive TME and enhancing cytotoxic T cell responses.[44] Hydrogels can also support synergistic chemo-immunotherapy, as demonstrated by DOX and R837 co-loaded PEG-based hydrogels that promote immunogenic cell death (ICD), dendritic cell maturation, and macrophage activation, resulting in effective inhibition of tumor growth and metastasis.[45]

Our study explored injectable hydrogels based on polyvinyl alcohol as base materials, but modified with carbohydrate or other gel-forming materials (alginate, chitosan, gelatine, galactomannan, Polyethylenimine), to assess their ability to modulate specific immune cell response. More importantly, a novel nano-adjuvant, based on Mn doped calcium phosphate, was incorporated into the hydrogel to activate innate immune pathways in infiltrated cells, targeting STING, TLRs, co-stimulatory molecules (e.g., CD86), and Th1 cytokines. By *in vitro* and *in vivo* studies we demonstrate that iHG is capable of selectively recruiting and activating immune cells. A critical question addressed was whether immune cells activated within this distant iHG could effectively target tumors at a remote site. Antitumor studies using a melanoma model demonstrated that this engineered iHG can robustly attract and activate immune cells outside the immunosuppressive tumor microenvironment (TME), eliciting a potent antitumor response without requiring neo-antigen vaccination or immune checkpoint blockade.

## 2 RESULTS AND DISCUSSION

### 2.1 Synthesis and characterization of immune-hydrogel

Different compositions of injectable hydrogels (iHGs) were developed using polyvinyl alcohol (PVA, 13.5% w/v) as the base matrix, supplemented with 0.1% (w/v) of alginate (iHG-ALG), chitosan (iHG-CS), galactomannan (iHG-GM), gelatin (iHG-GE), or polyethyleneimine (iHG-EI), and lyophilized under optimized conditions. To enhance immune activation, selected iHGs were further functionalized with STING agonist nanoparticles (iHG-STING), TLR9 agonist CpG (iHG-CpG), and immune checkpoint antibody (PD1). The morphology, porosity, and pore structure were analysed via SEM at 500X and 5000X magnifications (Figure 1B), revealing porous structures, with galactomannan-modified PVA gels exhibiting the largest pore size (∼5 µm). Micro-CT imaging confirmed a 3D porous architecture (Figure 1C), with a ∼40% porosity and an average pore volume of 0.1 cc/g (Supplementary Figure S1A, B), sufficient for the infiltration of immune cells (up to 20 µm in size). The rheological analysis demonstrated phase-angle independence of frequency, confirming gel-like and shear-thinning properties (Figure 1E, F). Immune modifiers (0.1%) did not significantly alter the gel’s rheological properties compared to the PVA matrix (Supplementary Figure S1C i–iv). The gels were physically robust and injectable through a 26-gauge syringe (Figure 1G, H). As shown in supplementary Figure S2, iHG gel in 1x PBS remained stable for up to 11 days at 24 °C, while it exhibited faster dissolution within 7 days when incubated at 37 °C The interconnected pore structure and antigen-loading capacity were evaluated using FITC-labelled ovalbumin. Cryo-sectioned fluorescence microscopy images (Figure 1I (i, ii)) showed uniform green fluorescence distribution across sections, confirming the interconnectedness of the pores for efficient antigen and immune adjuvant dispersion.

**Figure 1.**
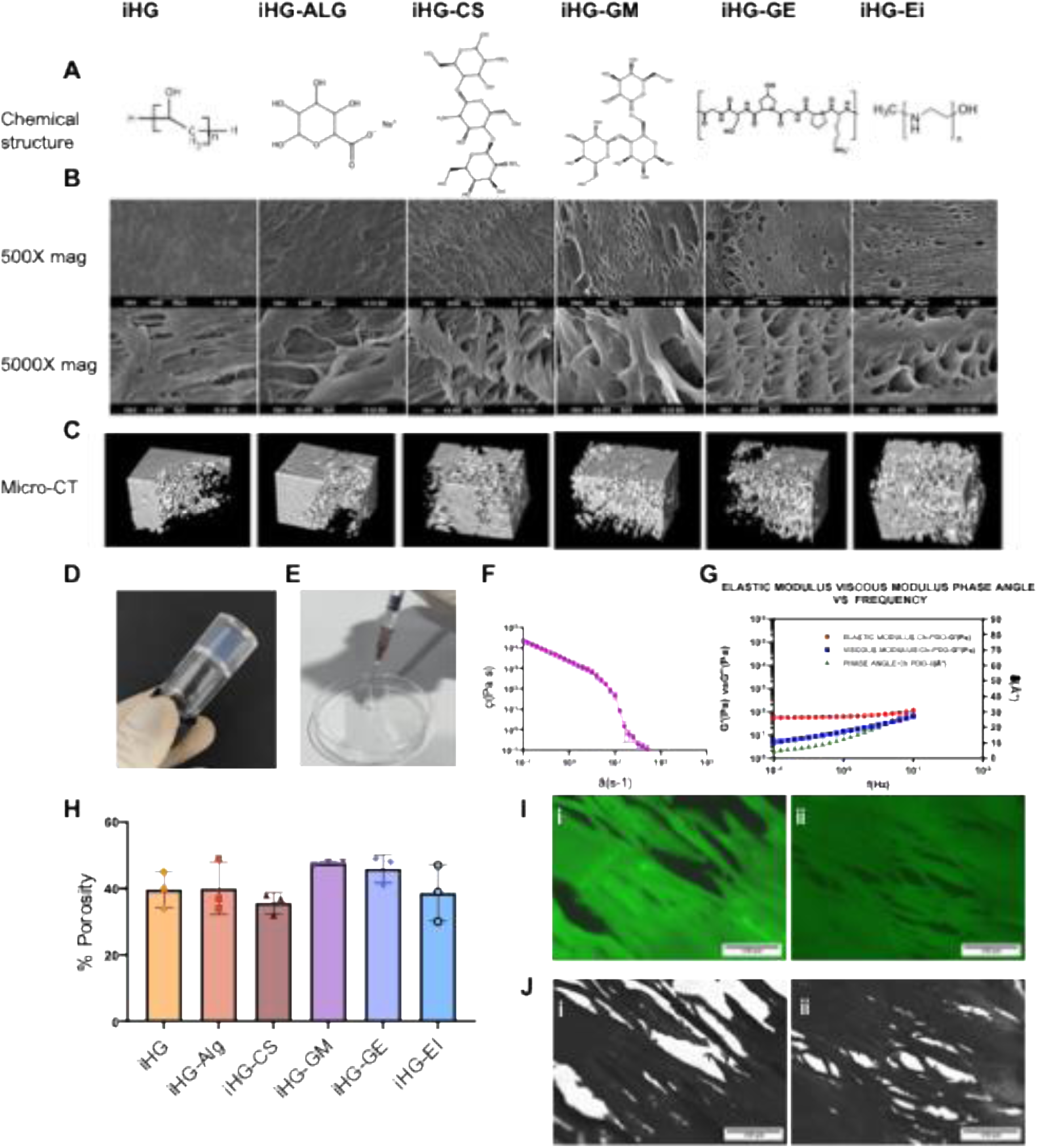
Characterisation of Immuno-hydrogel: (A) 2D chemical structure of different immune-modifiers used in the base 13.5% PVA (iHG) hydrogel with 0.1% (w/v) of alginate (iHG-ALG), chitosan (iHG-CS), galactomannan (iHG-GM), gelatin (iHG-GE), or polyethyleneimine (iHG-EI), (B) SEM images of surface pore structure of iHG gel matrix with different immune modifiers at 500x and 5000x magnification. (C) 3D pore structure analysis by Micro-CT imaging in contrast loaded iHG compositions, a cuboid volume section of 1.5mm x 1mm x 1.5mm size from each sample is compared (D) A representative image of freeze-thaw crosslinked PVA iHG (E) Injectability of cross-linked iHG through 26G needle. (F) Rheology analysis of iHG showing viscosity vs shear rate curve and (G) frequency sweep curve showing shear thinning and flow property of the gel. (H) Percentage porosity calculated from the micro-CT volume section (1.5×1×1.5mm) using ImageJ (BoneJ) shows an average of 40-50% porosity in 13.5% PVA gel matrix with different iHG compositions (I) Cryo-sectioned images of a lyophilized hydrogel loaded with FITC OVA albumin protein (200ug/ml), scale bar represents. (J) Phase contrast images of cryosection, 150 μm size. 2D chemical structures for gelatin, which was adapted from Milano et al. under a Creative Commons CC-BY license.[46]

### 2.2 *In vitro* biocompatibility and iHG characterization

To establish an artificial immune niche for immune cell infiltration, interaction, activation, and systemic dissemination, a biocompatible, porous, soft-tissue-mimicking microenvironment is essential. The biocompatibility of injectable hydrogels (iHGs) was evaluated using in vitro cultures of immune cells, including RAW 264.7 macrophages and JAWII dendritic cells. Cells were exposed to iHG formulations at concentrations ranging from 3.5 to 5.5 wt.% in culture medium for 24 hours (Figure 2A), assessing both cell viability and immune activation potential.

**Figure 2.**
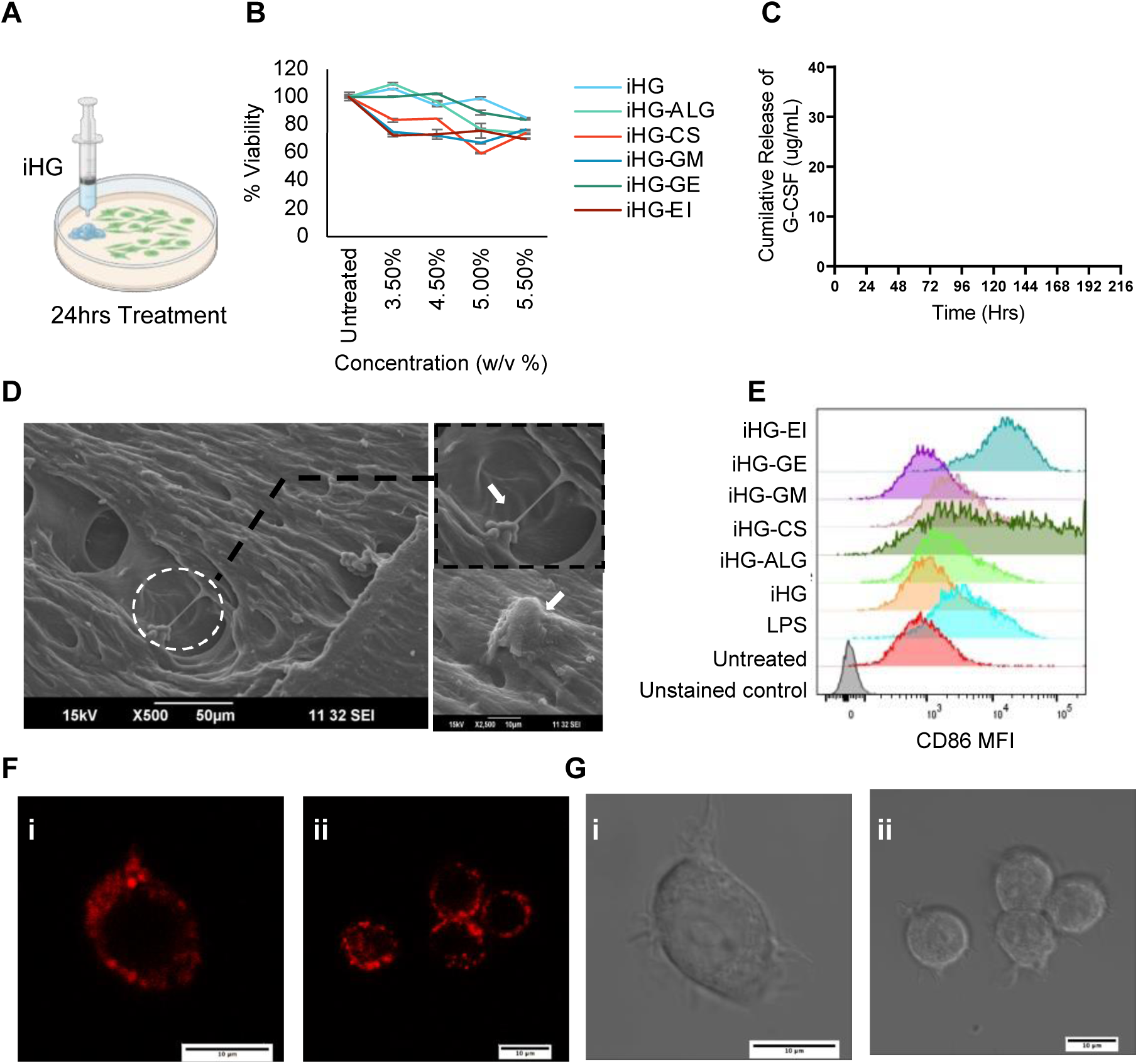
Invitro cell biocompatibility and activation. (A) shows schematics for in vitro cell culture treatment. (B) Viability of RAW 264.7 cells treated with different compositions of iHG in varying concentrations. Data represents mean ± SD for n=3 (C) *In vitro* cumulative release of loaded protein (G-CSF) up to 9 days in phosphate buffer at room temperature. Data represents mean ± SD for n = 3 (D) SEM images of RAW 264.7 cells growing in gel show the biocompatibility within the gel matrix. (E) Mean fluorescence intensity of CD86 expression in RAW 264.7 macrophage cells (F) RAW 264.7 macrophage cells showing intracellular uptake OVA-AF555 protein released from iHG after 24 hours of treatment under fluorescence and phase contrast microscopy.

Results (Figure 2B) demonstrated 70–80% viability of macrophages in iHG formulations containing 3–5.5wt/v% gel material. Among these, iHG-EI, incorporating 0.01% polyethyleneimine (PEI) with 13.5% polyvinyl alcohol (PVA) was optimized, as higher PEI concentrations were cytotoxic, probably due to cationic charge-mediated membrane disruption. Based on this, subsequent studies focused on the iHG-EI formulation with 0.01% PEI. To examine cell infiltration within the microporous structure, iHGs were incubated with RAW 264.7 macrophage cultures for 24 hours, followed by SEM imaging (Figure 2D). Cell clusters, exhibiting round and elongated morphologies, were observed within the gel-macro-pores and along their interior walls, confirming that the interconnected pore structure facilitated flow of medium and supports viable cell retention. To assess immune cell activation, CD86 expression, a co-stimulatory marker, was analysed (Figure 2E) in JAWs II dendritic cells post 24hrs treatment with iHGs. Both iHG-EI and iHG-CS formulations induced significant upregulation of CD86 compared to the positive control, LPS, demonstrating robust immune activation by these materials. The cationic charge of primary and secondary amines in PEI and chitosan likely enhanced interactions between cell phospholipids and the gel scaffold. These materials, known for their immune-activating properties, have been widely studied in nanoparticle formulations for antigen and RNA delivery in cancer vaccines, suggesting they may enhance antigen presentation and co-stimulation. Recent studies have highlighted the use of hydrogels and scaffolds to deliver biotherapeutics and modulate immune responses, such as in vaccines. [36,40,47]

The immune-activating microenvironment of the iHG can be engineered to retain antigens and release growth factors, stimulatory cytokines, or adjuvants that promote immune cell infiltration, activation, and maturation. These features may enhance the functional capabilities of antigen-presenting cells (APCs) that migrate into the hydrogel. By encountering neoantigens in the presence of adjuvants over an extended period, these APCs may exhibit improved antigen uptake, processing, and presentation, ultimately resulting in more effective immune priming and enhanced T cell infiltration. Careful selection of adjuvants that enhance antigen presentation is a critical consideration in designing effective immunotherapeutic platforms. o investigate the potential of the iHG for loading and controlled delivery of immune-stimulatory adjuvants, we examined the release kinetics of a model cytokine-adjuvant, G-CSF, over a 9-day period (Figure 2C). The iHG showed a cumulative release of approximately 30 µg G-CSF, with an initial burst within 24 hours, followed by a sustained release plateau after 72 hours. Furthermore, the antigen-loading capability of the iHG was evaluated using a model antigen, OVA-AF555, and its cellular uptake was assessed via fluorescence imaging. Confocal microscopy of RAW 264.7 macrophage exposed to OVA-AF555-loaded iHG for 24 hours revealed strong intracellular antigen uptake, as indicated by red fluorescence in the cell cytoplasm (Figure 2F). These findings indicate that the iHG can efficiently release the loaded neoantigen, and surrounding immune cells can uptake and process the antigens, enabling activation in a controlled, immune-favourable environment, away from the typically immunosuppressive tumor or peripheral microenvironment.

### 2.3 *In vivo* cell infiltration and cytokine profiling

Designing hydrogels to align with the migration, activation, and target specificity of the immune system involves creating intricate designs and properties crucial for effective cell modulation. These engineered hydrogels act as foundational elements in synthetic immune niches, 3D biomaterial structures that mimic natural lymph nodes. They provide a localized, alternative environment for immune cell interactions, expansion, and movement. Such synthetic immune niches have the potential to modulate the immune response against cancer and address challenges associated with immune-suppressed tumor-draining lymph nodes (TDLNs). Various humoral and cellular elements from both the innate and adaptive immune systems including neutrophils, macrophages, dendritic cells (DCs), lymphocytes, and the cytokines they secrete play crucial roles in eliciting an effective immune response. These factors interact with biomaterials in a spatio-temporal manner, thereby influencing and triggering the immune response. [29,48,49] To study the immune cell profile in such an artificial site created by different iHGs, we injected the hydrogel subcutaneously on the flank region of 12 weeks of healthy C57BL/6J mice and studied the infiltrated immune cells within the gel on day 5 as shown in Figure 3A.

**Figure 3.**
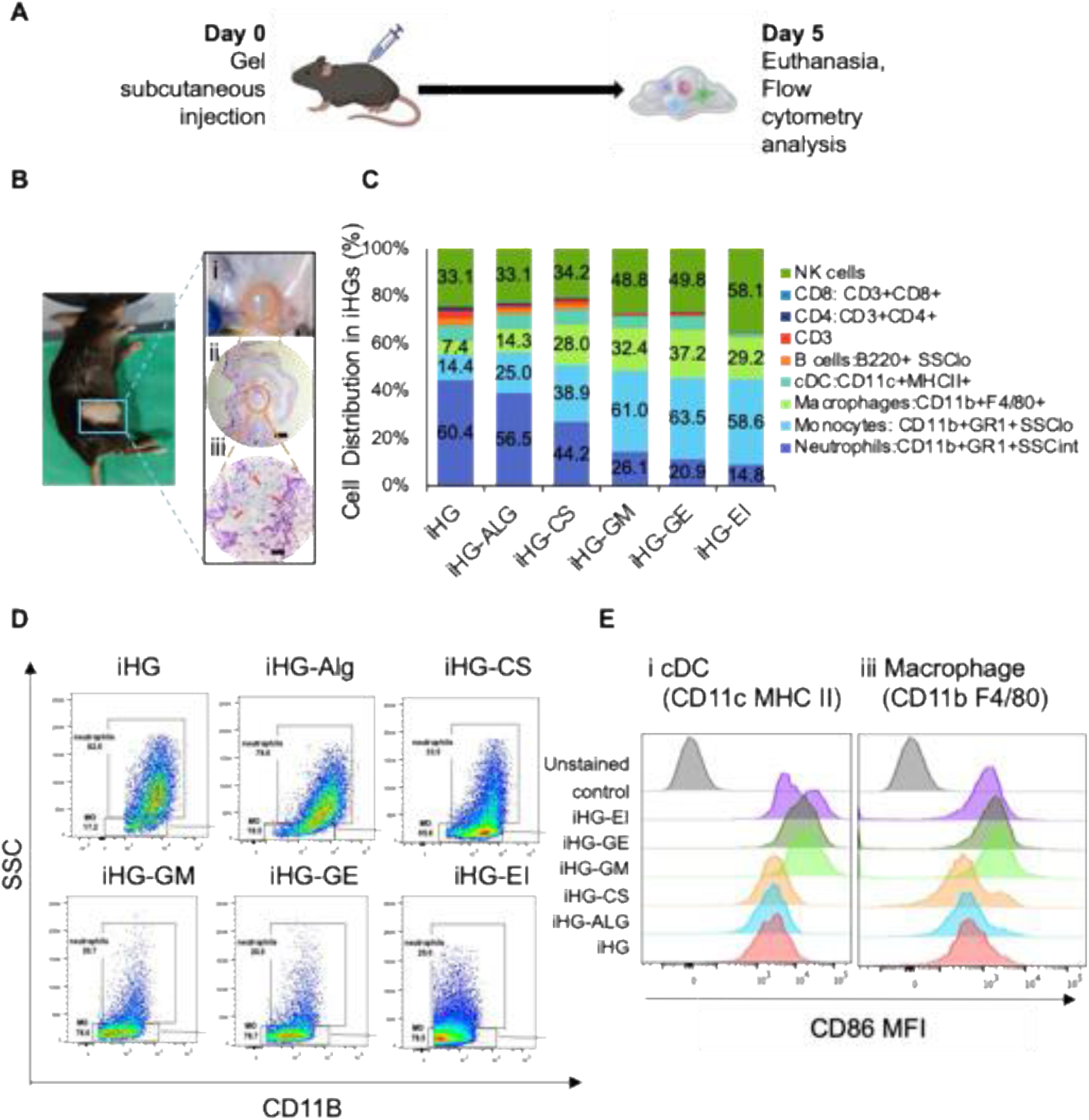
In vivo cell infiltration in iHGs. (A) shows schematics for in vivo gel injection in mice. (B) C57bl/6j with subcutaneous iHG injection: (i) shows the gel retrieved on day 5, (ii) H&E images of gel on day 5, scale bar 600μm (iii) Magnified image showing immune cell infiltration within the gel, scale bar 60μm (c) Flow cytometry analysis of different cell populations shows that varying composition can differentially attract specific cell types to the site of iHG injection, Data represents mean value, n=3 (D) Flow cytometry scatter plot SSC Vs CD11b showing the change in monocyte vs neutrophil population with different composition of iHG. (E) Shows the CD86 MFI activation of infiltrated (i) DCs (CD11c+ MHC-II+) and (ii)macrophages (CD11B+ F4/80+) cells.

Figure 3B shows that the gel depot formed subcutaneously in mice remained intact until day 5. The corresponding H&E images revealed cellular infiltration within the iHG (Figure 3B - i, ii, iii). We analysed the nature and frequency of infiltrated cells using flow cytometry (Figure 3C), which identified various immune cell types within iHGs, including NK cells, macrophages, monocytes, lymphocytes, and neutrophils. The frequency of these cell types varied depending on the gel composition. Compared to the base material PVA (iHG), chemically modified gels with different additives such as alginate, chitosan, galactomannan, gelatin, and PEI showed distinct shifts in the ratio of neutrophils to monocytes. Specifically, iHG-EI had a higher percentage of monocytes, while the base iHG had a high infiltration of neutrophils, as depicted in the scatter plot (Figure 3D). Additionally, there was detectable infiltration of CD3/CD4 and CD3/CD8 T cells in base PVA hydrogel and hydrogel modified with alginate or chitosan, whereas monocytes, macrophages and NK cells dominated in other compositions containing galactomannan, gelatin and PEI. This shows that iHGs may attract different types of immune cells based on their chemical composition. Further analysis of an important co-stimulatory marker, CD86, indicating the activation of dendritic cells (DCs) and macrophages, showed that iHG-EI, iHG-GE, and iHG-GM had higher expression than other gels (Figure 3E). This reflects their capacity to activate antigen-presenting cells (APCs) within the gel matrix. In the next step, we analysed the nature of cytokines released by the infiltrated cells within each iHG niche and blood plasma of the animals post-Day 5 of subcutaneous iHG injection. Figure 4A shows the experimental schematics for the detection of cytokine by flow cytometric bead analysis. The blood plasma cytokine profile in untreated healthy animals versus animals injected with different iHGs is shown in Figure 4B. While healthy animal only showed a detectable level of IL2, IFN γ and IL12, treatment with different iHGs showed an increase in pro-inflammatory cytokines such as IL6, IFN-γ, TNF-α and IL1-β and traceable amount of GMCSF, IL4, IL17a, IL2, IL4, IL10 response (iHG, iHG-AG, iHG-GE, iHG-EI). Plasma samples from iHG GM and iHG-CS showed high level of proinflammatory TNF and IFN-γ. To assess the cytokine profile of infiltrated cells and the gel matrix independently, we collected cell infiltrates and gel-associated fluid separately for multiplexed bead analysis using a comprehensive cytokine panel. Flow cytometric analysis (Figure 4C) of the infiltrated cells was performed, employing Uniform Manifold Approximation and Projection (UMAP) for dimensionality reduction and clustering to identify distinct cell populations within the gel matrix. The infiltrated cells were then lysed in the presence of protease and phosphatase inhibitors, and cytokine analysis of the lysates revealed a Th1-polarized pro-inflammatory milieu, characterized by elevated levels of IL-1β, TNF-α, and IL-6 (Figure 4D, F). This cytokine environment is conducive to an anti-tumor immune response, as efficient antigen presentation by dendritic cells or macrophages requires Th1 cytokines (signal 3) in the local microenvironment. Th1 cytokines, including interferon-gamma (IFN-γ), interleukin-2 (IL-2), and TNF-α, are pivotal in mediating anti-tumor immunity by enhancing interactions between immune and non-immune cells within the tumor microenvironment (TME). The presence of these cytokines within the gel matrix highlights its potential to establish a pro-inflammatory, immune-activating niche favourable for cancer immunotherapy. [50,51]

**Figure 4.**
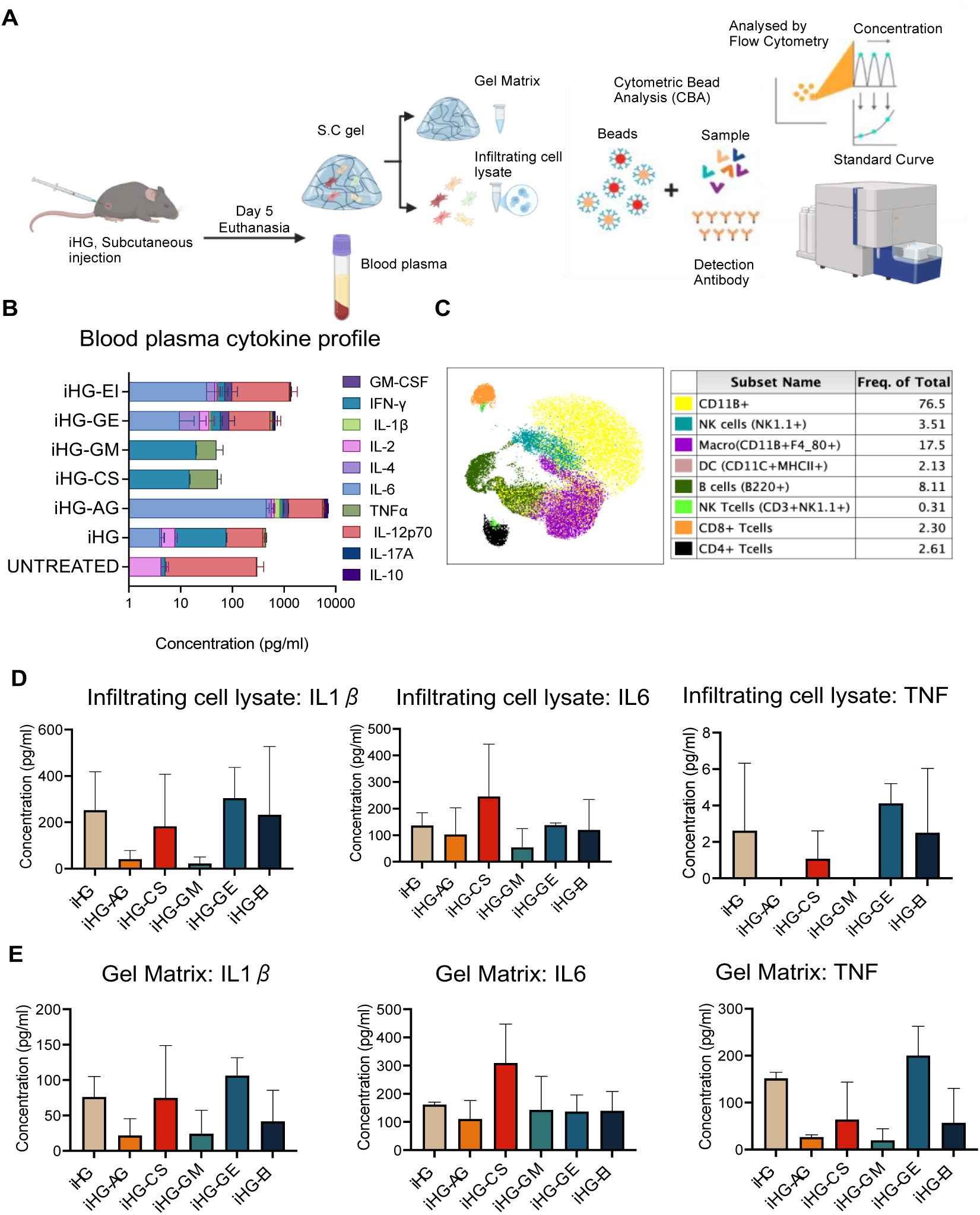
In vivo cytokine profiling of different iHGs. (A) Schematic showing experiment strategy and multiplexed CBA analysis of pro-inflammatory and anti-inflammatory cytokine 5 days after injection with iHGs (B) cytokine profile of blood plasma post-injection. (C) UMAP clustering of cells infiltrating in subcutaneously injected iHG gel after 5 days showing population clusters of lymphocytes (CD4+, CD8+, B220+, NK1.1+) and myeloid cell population (CD11b+, CD11c+, F4/80+). (D) Concentration (pg/ml) of pro-inflammatory cytokine IL1-*β*, TNFα, and IL6 in cell lysate prepared from the infiltrated cells in a gel matrix. (E) shows the cytokine level of pro-inflammatory cytokine IL1-*β*, TNFα, and IL6 in the gel matrix collected on day 5 post subcutaneous injection, data represents mean ± SD for n=3.

### 2.4 *In vivo* antitumor study using iHGs

To investigated the potential of immune hydrogels (iHGs) to enact an anti-tumor immune response by facilitating immune cells to assemble at the site of injection, away from the typical immune-compromised TME or peripheral immune organs. In a murine melanoma model using C57BL/6J mice, we evaluated the therapeutic efficacy of injectable hydrogels (iHGs) as an antigen-free immunotherapy. As illustrated in the schematic (Figure 5A), mice received a single subcutaneous injection of iHGs on day 8 following tumor inoculation. For the combination therapy group and anti-PD1 alone control group, anti-PD-1 antibody was administered intraperitoneally on days 10 and 15. Tumor progression was monitored up to day 21 post-inoculation. Unmodified iHG was used as a material placebo and four different modified iHG compositions were tested as: i) a novel adjuvant nanoparticle, (Ca,Mn)PO4, activating stimulator of interferon genes (STING), incorporated within the iHG matrix forming iHG-STING gel ii) cationically modified iHG-STING gel using PEI as polymeric adjuvant (iHG-STING-EI), iii) a standard adjuvant CpG activating TLR9 forming iHG-STING-CpG and iv) iHG-STING-CpG with clinically approved immune checkpoint blockade antibody, anti-PD1.

**Figure 5.**
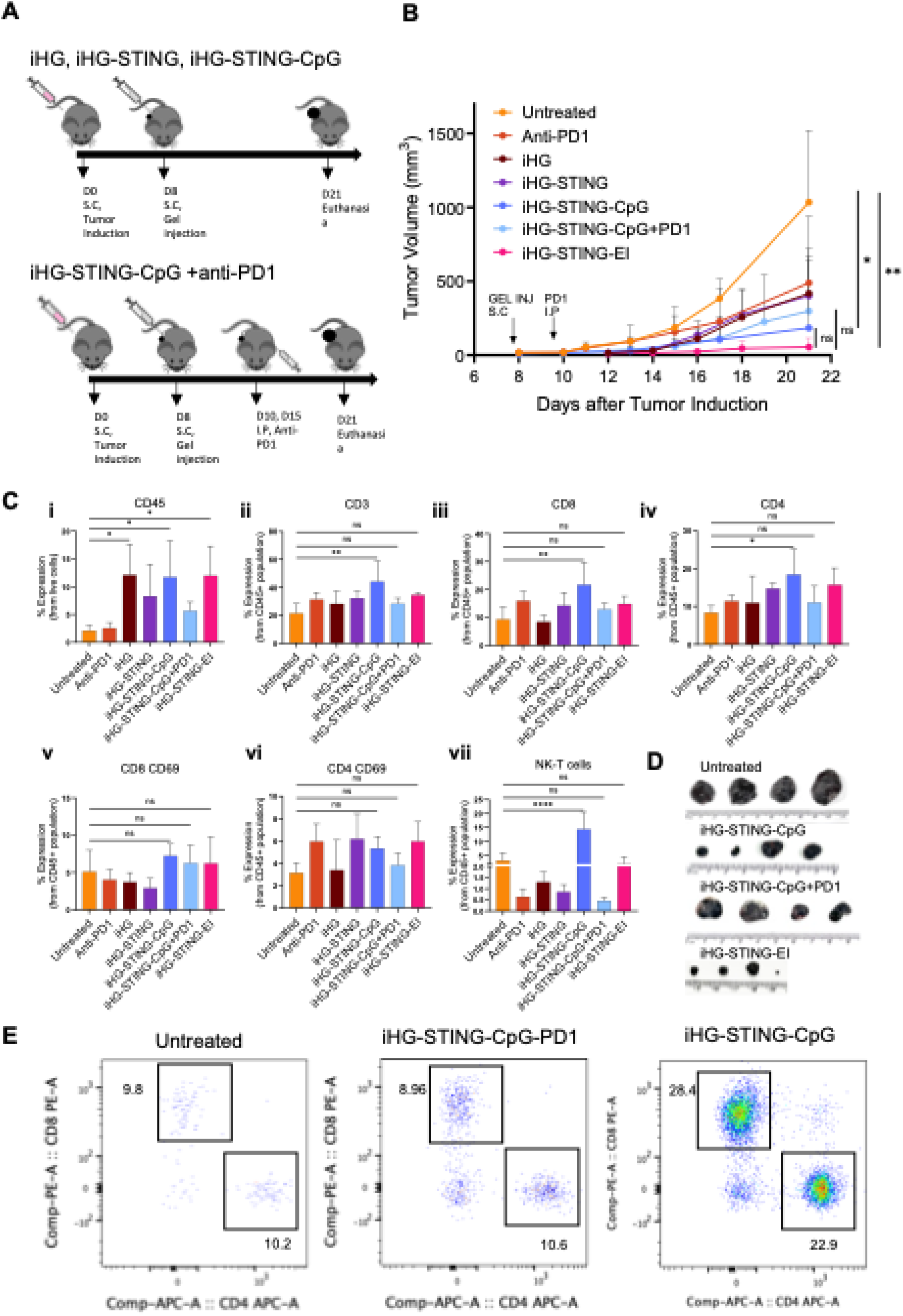
iHG-combinatorial anti-tumor immunity with ICB therapy. **(A)** shows schematics for anti-tumor study in mice subcutaneous melanoma model for iHG, iHG-STING, iHG-STING-CpG, and iHG-STING +anti-PD1 treatment groups. (B) Tumor growth curve up to 21 days post B16F10 tumor induction for different treatment groups. (C) flowcytometry analysis of infiltrated immune cells in tumor post 14 days of treatment shows an increase in t cells (CD4, CD8, NKT) and its activation in iHG-STING-CpG groups. (D) Tumor images of untreated, iHG-STING-CpG, iHG-STING-CpG+PD1 and iHG-STING-EI post euthanasia on day 21. (E) shows the scatter plot of CD4 vs CD8 population from CD45+ cells in tumor. Data are presented as mean ± SD (n = 4). Statistical comparison in tumor growth in (B) (day 21) was performed using ordinary one-way ANOVA with Tukey’s multiple-comparisons test. Statistical comparisons in flow cytometry analysis (C) performed using one-way ANOVA with Dunnett’s Multiple Comparison against untreated control group.

The cGAS–STING pathway mediates innate immune activation, particularly in dendritic cells and macrophages. Upon activation, this pathway initiates a transcriptional program that leads to the production of type I interferons (IFNs) and proinflammatory cytokines.[52,53] Type I IFNs subsequently drive the expression of interferon-stimulated genes (ISGs), which enhance antigen presentation, promote dendritic cell maturation, upregulate MHC class I expression, and augment the cytotoxic functions of natural killer (NK) cells and CD8⁺ T cells.[52,54] Collectively, these effects enable the cGAS–STING axis to act as a critical bridge between innate and adaptive immunity. Research by Wang et al. demonstrated that Mn^2+^ ions enhance the sensitivity of the cGAS-STING pathway.[55] Subsequent studies by various groups have indicated that nanoparticles or nano complexes containing Mn^2+^ can stimulate tumor-infiltrating dendritic cells (DCs), which in turn activate CD8+ T cells and boost the activity of natural killer (NK) cells against tumors.[56–58] Accordingly, we have synthesized a novel Calcium-Manganese phosphate (Ca,Mn)P nanoparticle adjuvant capable of effectively activating STING, IFN-β pathway and enhanced co-stimulatory expression (CD40, CD 86, and MHC-II) (supplementary Figures S3) and the same was loaded into the iHG to form a potent innate activating iHG-STING gel. The uniform distribution of these NPs in the gel matrix and activation of STING in dendritic cells tested *in vitro* is shown in Supplementary Figure S3. Further, we asked the question whether the cationic modification of iHG-STING by way of modifying with PEI will enhance the co-stimulation or addition of a clinically relevant and approved TLR agonist (TLR9) in iHG-STING will improve the anti-tumor-response. In another aspect, while CD8+ T cell-mediated cancer clearance is crucial, it is often undermined by the interaction between check point inhibitory molecules like PD-1 and PD-L1. This interaction serves as a brake to prevent excessive T cell activation under normal conditions but is exploited by tumor cells to evade immune detection. Immune checkpoint inhibitors (ICIs) have revolutionized cancer treatment in recent times by removing these inhibitory breaks, thereby boosting T-cell activity against tumors[15,16]. However, in the current clinical scenario, only < 25% of cancer patients respond to ICI immunotherapies, likely due to inadequate immune activation posed by compromised TME. As a result, combining anti-PD-1 therapies with iHG-type engineered immune niche therapeutics is a potential combinatorial approach and we investigated the same in the melanoma model.

The anti-tumor therapeutic effect of all the iHG compositions is presented in Figure 5B. Interestingly, the PVA-based iHG platform alone showed an impact on tumor growth comparable to that of anti-PD1 antibodies. Enhanced tumor suppression was observed with iHG modified with STING-activating (Ca,Mn)PO₄ nanoparticles (iHG-STING) and further improved with co-loading of the TLR9 agonist CpG (iHG-STING-CpG), figure 5D shows the images of the tumor collected on day 21 post euthanasia and supplementary figure S4 shows the tumor images of all treatment groups. Notably, combining anti-PD1 therapy with the treatment of iHG-STING-CpG did not yield any significant improvement. Interestingly, the cationically modified iHG-STING-EI, exhibited almost the same tumor control compared to both iHG-STING-CpG or its combination with anti-PD1. Statistical analysis using one-way ANOVA showed significant tumor volume reduction in both iHG-STING-CpG and iHG-STING-EI groups compared to the untreated control group. This indicates that suitably designed iHGs have the potential to enhance the antitumor immunity.

The intra-tumoral cell response analysis using flowcytometry revealed significant enhancement in the infiltration of CD45^+^ cells to the tumor, in iHG, iHG-STING-CpG and iHG-STING-EI implanted animal groups (Figure 5C (i)), clearly indicating enhanced immune-cell response that correlates with the tumor growth control. More importantly, the lymphocyte fraction showed elevated levels of CD3, CD4, CD8 T cells and NK T cells in iHG-STING-CpG and iHG-STING-EI groups(Figure 5C (ii-vii). Figure 5D presents a representative scatter plot illustrating the enhanced infiltration of CD4⁺ and CD8⁺ T cells into the tumor microenvironment following treatment. CD8⁺ T cells, recognized as the primary effector cells of the adaptive immune response, showed significant recruitment. This outcome is particularly promising, as it was achieved solely through the administration of the engineered biomaterial gel system, without requiring antigen-specific vaccination.

These findings highlight the therapeutic potential of immunologically modified iHGs as a novel class of immunotherapeutics. By creating a supportive immune microenvironment that facilitates immune cell infiltration, activation, and dissemination, iHGs enable effective anti-tumor responses, even without any neo-antigen vaccination and in the presence of pre-existing systemic immune suppression. This approach represents a paradigm shift in the use of novel biomaterials for cancer immunotherapy, offering a strategy to bypass immune suppression in the tumor microenvironment.

## 3 Conclusion

In conclusion, our study demonstrates the potential of injectable biocompatible polymer hydrogels, modified with immune modulators, as an innovative platform for attracting and activating immune cells at locations distant from the immune-suppressed tumor microenvironment or peripheral organs. These hydrogels effectively mobilize immune cells to combat tumors at a remote site. We validated the injectable and biocompatible properties of the porous hydrogel matrix, its ability to load and release therapeutic proteins, and its capacity to incorporate novel immune-modulating adjuvants. The hydrogels activated innate immune pathways, such as STING and TLR, and enhanced CD86 co-stimulatory molecule expression, promoting anti-cancer immune response.

In vivo studies in healthy and melanoma models showed that the immune-hydrogel attracted diverse class of immune cells, maintained a Th1-polarized cytokine milieu, and enhanced CD45⁺ immune cell infiltration, including CD4⁺ and CD8⁺ T cells and NK cells into the tumor microenvironment. Notably, these results were achieved without neo-antigen vaccination or immune checkpoint blockade. Moreover, the possibility of repeated administration of biodegradable iHGs for periodic recruitment of immune cells and their activation to continuously fight cancer is a significant milestone, especially from the perspective of metabolic exhaustion faced by T cells in the current immunotherapeutics. In summary, our findings position iHG systems as a promising cancer immunotherapy candidate, warranting further studies to explore their safety, efficacy, and applicability across diverse tumor types and immune settings for potential clinical translation.

## 4 Experimental Section

### Materials

Polyvinyl alcohol (PVA), Sodium Alginate from brown algae, Chitosan, Gelatin from porcine skin, Locust bean gum from Ceratonia siliqua seeds (Galactomannan), and polyethylene imine (PEI) were purchased from Sigma-Aldrich. The 2D structure of gelatin was adapted from an open-access article published by Milano et al.under a Creative Commons CC-BY license, while other structures were sourced from PubChem.[46] Raw 264.7 (mouse macrophage cell lines), JAWS (immature dendritic cell lines), and B16F10 (murine melanoma cell line) were acquired from ATCC. Dulbecco’s Modified Eagle Medium (DMEM) high glucose media, Minimum Essential Medium Eagle (MEM-αmodification), Fetal Bovine Serum (FBS), and antibiotic-pen strep were bought from HiMedia. Alamar Blue, a cell viability reagent, was purchased from Invitrogen. Tissue Culture plate Sterile PS 48 wells and 12 wells were bought from Tarsons. 70µm cell strainer was procured from Falcon. CD16/32, CD45-APCR700, CD3 FITC, CD4 APC, CD8 Percp Cy5.5, CD11b FITC, CD11c BV421, MHC II PE, B220 BV610, Ly6G BV711, GR1 PE, CD86 PE Cy7, CD49b BV510, NK1.1 BV711, CD69 PE-Cy7 antibodies were purchased from BD Biosciences.

### Hydrogel preparation and characterization

13.5% PVA and 13.6% PVA-Alginate, PVA-Chitosan, PVA-Galactomannan, PVA-Gelatin, and PVA-PEI were prepared using the freeze-thaw method. All the polymers were weighed, and the polymer solutions were made by adding polymers in addition to milli-Q and constant stirring using a magnetic stirrer (Thermo-Fisher Super Nova Multi-position Digital Stirring Hotplates Stirrer). The stirred polymer solutions were then mixed based on their specific combinations, and the mixtures were kept in −80^ο^C overnight and thawed at room temperature for 1 hour. The porosity of the obtained hydrogels was analysed using a Scanning Electron Microscope (JEOL JSM-64901A). Lyophilized samples were taken for imaging, and prior to imaging, lyophilized hydrogels were sputter-coated with gold and visualized at 20 kV. The three-dimensional porosity of the hydrogel was imaged using Micro-CT (MI Labs BV). The lyophilized samples were visualized at 50 kV tube voltage and 0.24 mA tube current. The porosity of the hydrogels was qualitatively analysed using Imalytics software and quantitatively analysed using ImageJ software. Rheological characterization of hydrogels was done using Malvern Kinexus Pro, UK equipped with a parallel plate of diameter 20mm and 0.5mm gap at 37^ο^C. Viscoelastic properties of the hydrogels were analysed by performing frequency sweep and flow curve analysis. The three main parameters storage modulus (G’), Loss modulus (G’’), and loss tangent (G’’/G’) were obtained from a frequency sweep carried out from 1×10^-1^ to 1×10^2^Hz, which reflected the hydrogel behaviour at different time scales. Flow curve analysis was done to provide insights into the flow property and viscosity of hydrogels with shear rates ranging from 1×10^-1^ to 1×10^3^ s^-1^.

#### Cell culture

Three types of cell lines were used for this study, namely Raw 264.7, JAWS, and B16F10. Raw 264.7 and B16F10 cells were cultured in DMEM media with (10% FBS and 1% antibiotic-pen strep). JAW cells were cultured in α-MEM with (20% FBS, 1% antibiotic-pen strep, 5µg/ml GM-CSF, and sodium pyruvate). All cells were maintained at 37^°^C and 5% Co_2_.

### Invitro cell biocompatibility

The toxicity of hydrogels was analysed using an Alamar Blue cell viability assay. Raw 264.7 cells (1.5×10^4^) were seeded into a 48-well plate along with DMEM media and were incubated for 24 hr at 37^ο^C. Then, the cells were treated with different concentrations of hydrogel, followed by the next 24 hr incubation. After the incubation time, 10% of the total volume of the well of Alamar blue reagent was added into all the wells and kept for 4 hr incubation. After the final incubation, fluorescence values were obtained from a microplate reader, which is used for calculating cell viability. To image cells growing within the matrix, 13.5% gel was cross-linked and lyophilized. 1×10^5^ Raw 264.7 cells were seeded onto the lyophilized gel in a 12-well plate with DMEM complete media. After 24 hours, the lyophilized gel was sectioned and fixed in 4% paraformaldehyde (PFA). The section was analysed by Scanning Electron Microscope (JEOL JSM-64901A) after alcohol dehydration and gold sputtering.

#### Confocal uptake

OVA-AF555-loaded iHG was incubated with RAW 264.7 macrophage cells to assess cellular uptake. Briefly, 100 µL of OVA-AF555-loaded hydrogel was added to RAW 264.7 cells cultured in a 12-well glass-bottom plate at a density of 1 × 10⁵ cells per well. The cells were incubated with the hydrogel at 37°C in complete DMEM media for 24 hours to allow for uptake. After incubation, the cells were washed with PBS to remove any unbound hydrogel. Cellular uptake of the OVA-AF555-loaded hydrogel was then assessed using a Leica confocal microscope, with fluorescence detection of AF555 (excitation: 555 nm, emission: 565–585 nm) alongside phase contrast imaging to visualize cell morphology.

#### *In Vitro* Peptide Release

iHG was mixed with G-CSF (filgrastim, 30 µg) and cross-linked using the freeze-thaw method. The resulting cytokine-loaded gels were incubated at room temperature in 1 mL of release buffer (1× PBS). At the indicated time points, the release buffer was collected and replaced with 1 mL of fresh buffer. The amount of released cytokine was quantified using a pre-coated Human G-CSF Instant ELISA (Invitrogen). Briefly, test samples were diluted 1:4 and added to the capture antibody-coated wells along with Biotin-Conjugate (anti-G-CSF polyclonal antibody) and standards. The plate was incubated at room temperature (18-25°C) for 3 hours, then washed 6 times with 400 µL of 1× wash buffer. Subsequently, 100 µL of TMB Substrate Solution was added to all wells and incubated at room temperature for 10 minutes until colour developed in the top standard. Finally, 100 µL of Stop Solution was added to all wells, and colour intensity was measured at 450 nm. A standard curve was generated by plotting the mean absorbance for each standard concentration, and a best-fit curve was drawn through the data points. The concentration of G-CSF in the test samples was determined by extrapolating the absorbance values from the standard curve and multiplying by the dilution factor.

### JAWS cell activation study

In vitro CD86 expression of JAW cells exposed to different hydrogel combinations was analysed by FACS. JAWS cells (1×10^5^) were seeded into a 12-well plate along with α-MEM media and were incubated for 24 hours at 37^ο^C. Then, the cells were treated with two concentrations of the different hydrogels, followed by incubation for 24 hours. After the incubation time, the treated media was removed, and the cells were detached from the wells using 0.5% trypsin. After washing the cells with phosphate buffer solution (PBS) by centrifuging at 500 rpm for 5 minutes, the cells were incubated in blocking solution (10% BSA) for 30 minutes. Then, before the addition of CD86 antibodies, cells were again washed using PBS and followed by FACS.

### In Vivo Immune Cell Infiltration

All in vivo experiments were conducted following the guidelines of the Committee for the Purpose of Control and Supervision of Experiments on Animals (CPCSEA), Government of India. The study protocol was reviewed and approved by the Institutional Animal Ethics Committee (IAEC) at Amrita Institute of Medical Sciences, Kochi, Kerala, India (Approval No. IAEC/2021/1/4).

Briefly, the study was conducted C57BL/6 (n=3, per treatment group) mice were subcutaneously injected with 200 µL of each hydrogel combination. After 5 days, mice were euthanized, and the hydrogels were retrieved. Blood was collected via cardiac puncture into EDTA-coated vials, and plasma was separated by centrifugation at 4500 rpm for 20 minutes at 4°C. The plasma was stored at −20°C for later multiplexed cytokine analysis. The hydrogels were processed by passing them through a 40 µm cell strainer to separate cells from other debris. The cells were washed with PBS by centrifugation at 500 rpm for 5 minutes. The gel matrix supernatant, containing soluble cytokines, was stored at −20°C for further analysis. The collected cells were aliquoted for flow cytometry analysis and for preparation of cell lysates using Cell Extraction Reagent (Invitrogen) supplemented with PMSF and protease inhibitor cocktail (PIC). The cell lysates were stored at −20°C for subsequent cytokine analysis. at −20^0^C for cytokine analysis.

### Cytokine analysis

Cytokine levels in mouse plasma or cell culture supernatants were quantified using a Mouse Flex Set (BD Biosciences) for GM-CSF, IL-4, IL-17A, IL-2, IL-10, IFN-γ, IL-12, TNF-α, and IL-1β by Cytometric Bead Analysis (CBA). Standard cytokine concentrations were prepared using serial dilutions, and 50 µL of sample was incubated with 50 µL of specific bead mixture for 30 minutes at room temperature. After washing with PBS, PE-conjugated detection antibodies were added and incubated for an additional 30 minutes. Following washing, samples were resuspended in 300 µL staining buffer and analysed by flow cytometry on a BD FACS Lyrics. Data were collected with at least 10,000 events per sample, and cytokine concentrations were determined by comparing sample median fluorescence intensity (MFI) to a standard curve, adjusted for dilution factors.

#### Tumor study

C57/BL6 (12-14 weeks, 18-21g, n=4 per treatment group) were subcutaneously injected in the right flank with B16F10 cells (1×10^5^) on day 0 to induce the tumor. By Day 7, once tumors became palpable, mice were randomized into different treatment groups. On day 8, mice received subcutaneous injections of various hydrogel formulations, including iHG, iHG-STING, iHG-STING-CpG, and iHG-STING-EI. In addition, mice in the iHG-STING-CpG +PD1 group and a separate anti-PD-1 control group received intraperitoneal injections of anti-PD-1 antibody (12 mg/kg body weight) on Days 10 and 15. Tumor growth was monitored until Day 21. Mice were then euthanized, and tumors were harvested for flow cytometry analysis to evaluate the immune cell populations within the tumor microenvironment.

### Flow cytometry

Tumor tissues were processed and stained with fluorophore-conjugated primary antibodies for flow cytometry analysis. Briefly, tumors were washed with 1× PBS and minced into small fragments (3–4 mm). Enzymatic digestion was performed using a solution of collagenase and DNase I in 1× PBS, followed by incubation in a shaking incubator at 37 °C for 45 minutes. The resulting cell suspension was passed through a 70 µm cell strainer to remove debris and extracellular matrix. Red blood cells were lysed using RBC lysis buffer, and the remaining cells were washed with PBS by centrifugation at 500 × g for 5 minutes. For antibody staining, 1 × 10⁶ cells per sample were first incubated with mouse Fc block (anti-CD16/32) to prevent nonspecific binding, followed by viability staining. Cells were then washed and incubated with a cocktail of fluorophore-conjugated antibodies targeting mouse CD45, CD3, CD4, CD8, CD69, NK1.1, and CD49b for 20 minutes at room temperature. After staining, cells were washed to remove excess antibody and analysed using a flow cytometer.

### Statistical Analysis

All data are presented as mean ± standard deviation (SD). Statistical analyses were performed using GraphPad Prism version 9 (GraphPad Software, San Diego, CA, USA). For the anti-tumor study, a sample size of n = 4 per group was used. One-way analysis of variance (ANOVA) was conducted to compare tumor growth on day 21, followed by Tukey’s multiple comparison post hoc test to assess differences between groups. Comparisons of infiltrated immune cell populations were performed using Dunnett’s multiple comparison post hoc test against the untreated control group. A p-value ≤ 0.05 was considered statistically significant.

## Supporting Information

Supporting Information is available from the Wiley Online Library or the author.

## Acknowledgments

The authors gratefully acknowledge the Department of Biotechnology (DBT) and Swiss National Science Foundation under the Indo-Switzerland Joint Research Programme (ISRJP) project titled “Combinatorial activation of anti-tumor immunity using innovative multi-targeted nano Immunotherapeutics”(BT/IN/Swiss/56/MK/2018-19) for the financial support. The authors thank the contribution of Mr. Sajin P Ravi, Mr. Dennis Mathew, and Dr. Siddharamana Gowd for SEM and Micro CT imaging and Central Lab Animal Facility, AIMS, Kochi, for conducting in vivo animal experiments. The authors thank Amrita Vishwa Vidyapeetham for all infrastructure support.

## Supporting Information

**Figure S1.**
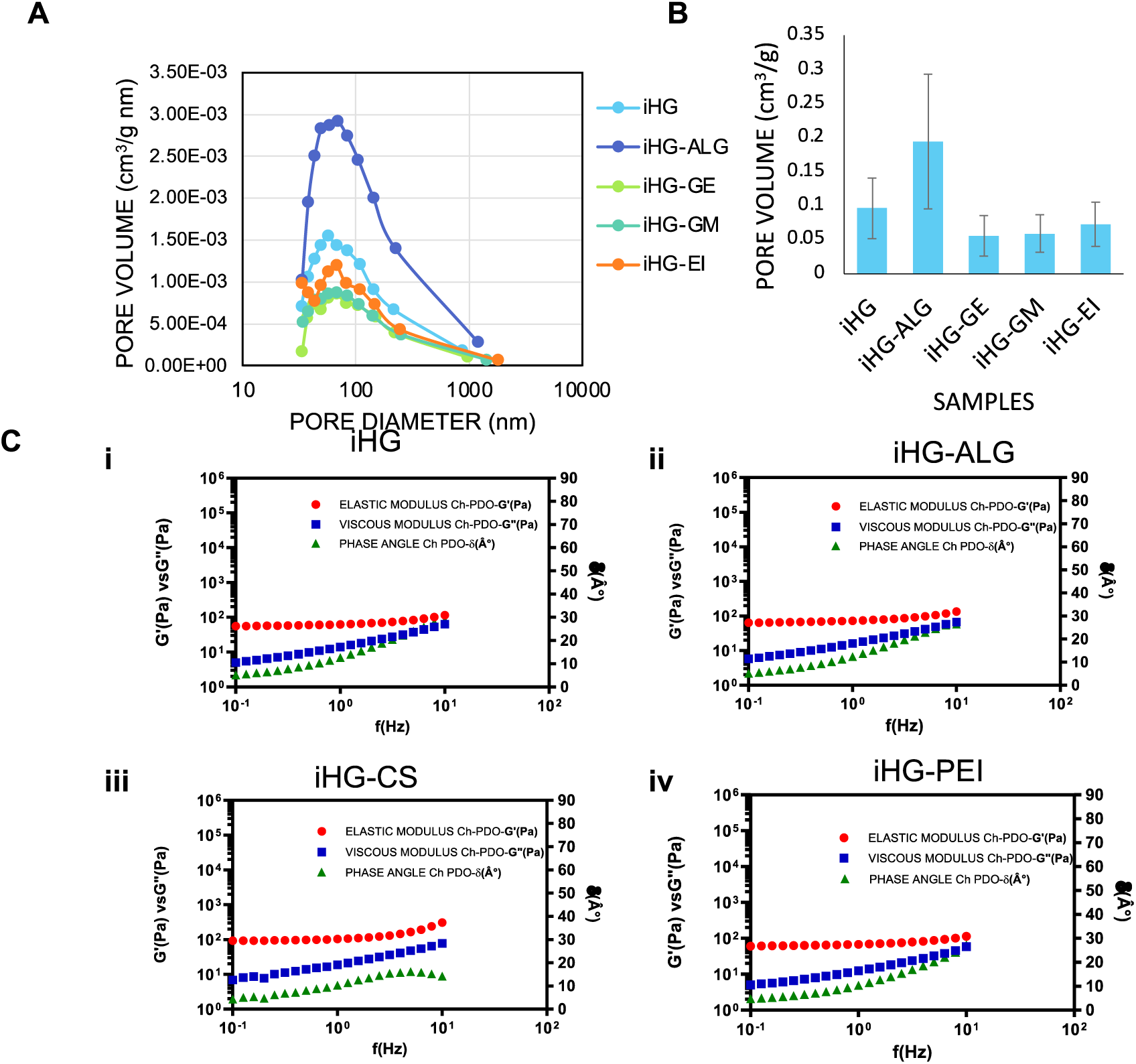
Characterisation of Immunohydrogel (iHG) (A) BJH pore size distribution curves for iHGs with varying compositions, highlighting differences in pore size and volume across samples. (B) Bar graph showing the average pore volume of approximately 0.1 cm^3^/g for iHGs with different compositions, derived from BET analysis for iHGs with different compositions. (C) Rheological analysis of gels with varying compositions (i–iv). Frequency sweep curves demonstrate the shear-thinning behavior and consistent flow properties of the gels.

**Figure S2.**
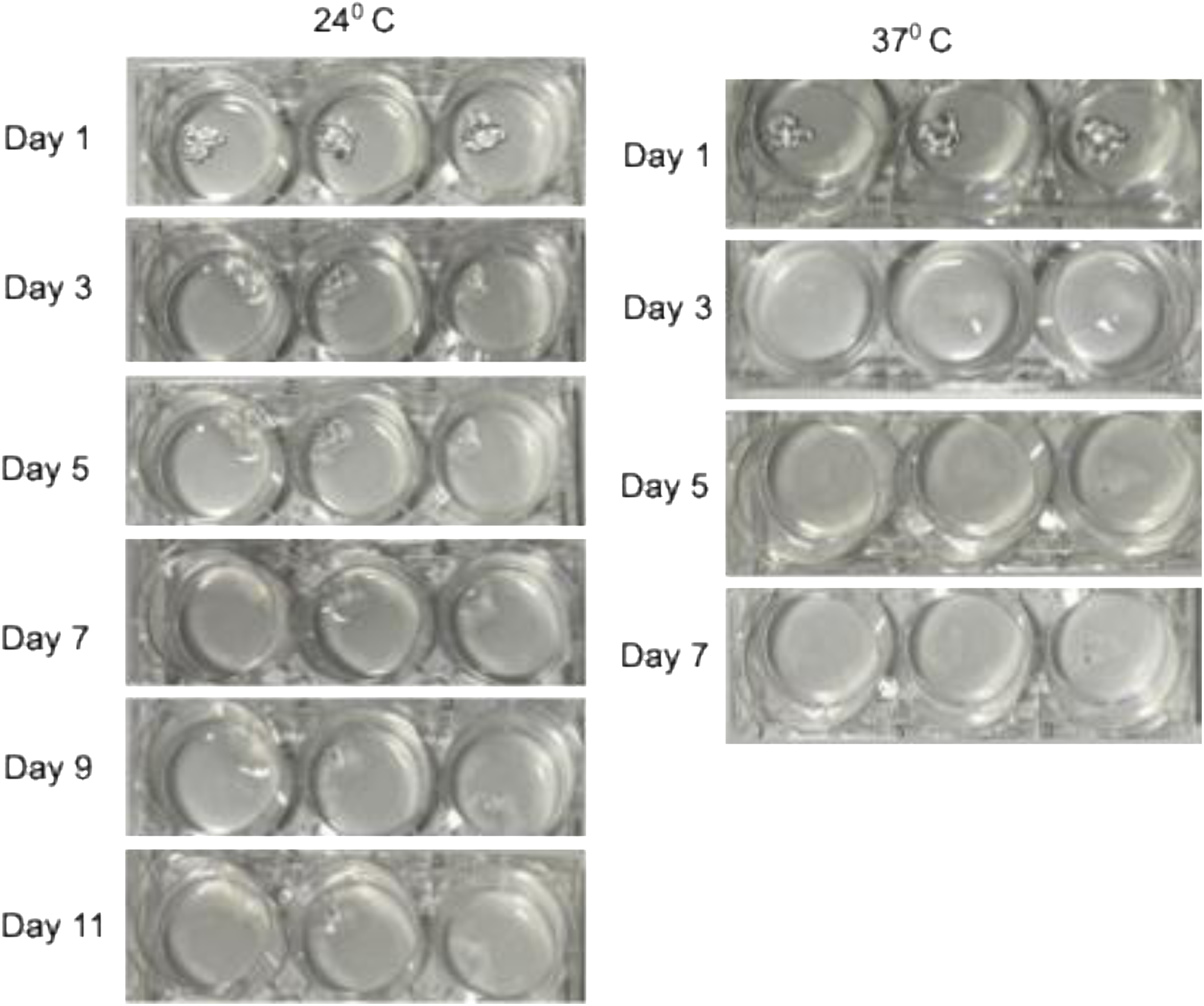
In vitro degradation study of iHG gel. (A) Representative images showing gradual degradation of the gel over 11 days in 1× PBS at 24 °C, indicating slow dissolution. (B) Faster degradation observed within 7 days in 1× PBS at 37 °C.

**Figure S3.**
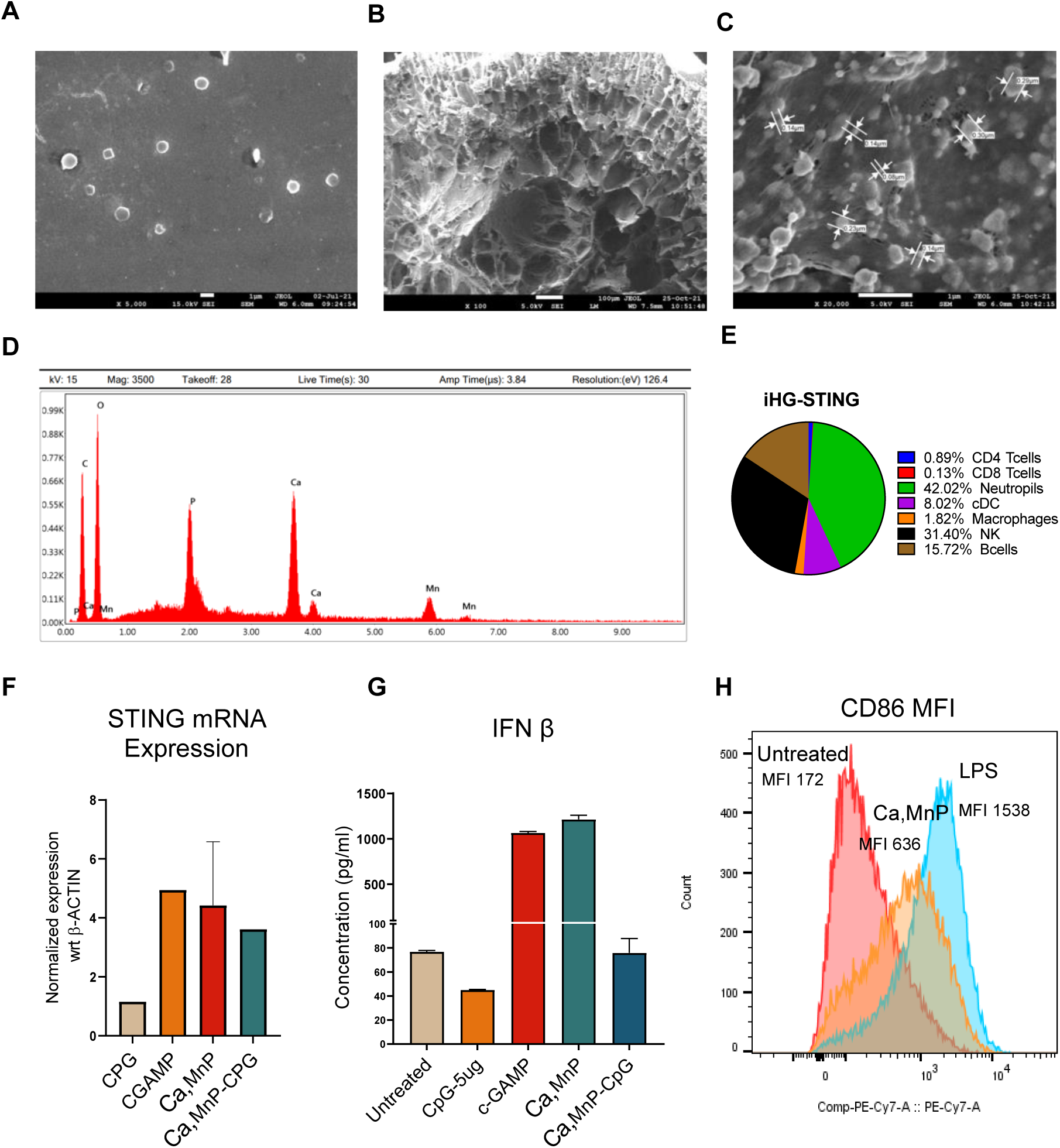
iHG STING characterization. (A) SEM images of (Ca, Mn)P nanoparticles showing multidispersered nanoparticles with size range from 1000μm to 100nM. (B) SEM images (100x) of a lyophilized section of iHG loaded with STING activating nanoparticles. (C) High magnified images of (Ca, Mn)P nanoparticle loaded within iHG gel. (D) EDAX spectrum of loaded nanoparticle showing a peak of calcium and manganese within iHG. (E) Immune cell infiltration in iHG-STING gel post 5 days after subcutaneous implantation in C57BL/6J mouse, showing a high infiltration of neutrophils, NK cells, B cells, and DC. (F)shows the increase in the mRNA expression level of STING after 24 hrs treatment 100μM (Ca, Mn)P and (Ca, Mn)P+CpG in the JAWS II DC cell line. (G) Shows the release of type I interferon (IFN-*β*) by STING activation by JAWS II DC cell line with the treatment of 100μM (Ca, Mn)P and (Ca, Mn)P+CpG. (H) Increase in CD86 activation (MFI) in JAWS II DC with treatment with 100μM (Ca, Mn)P nanoparticle for 24 hours.

**Figure S4.**
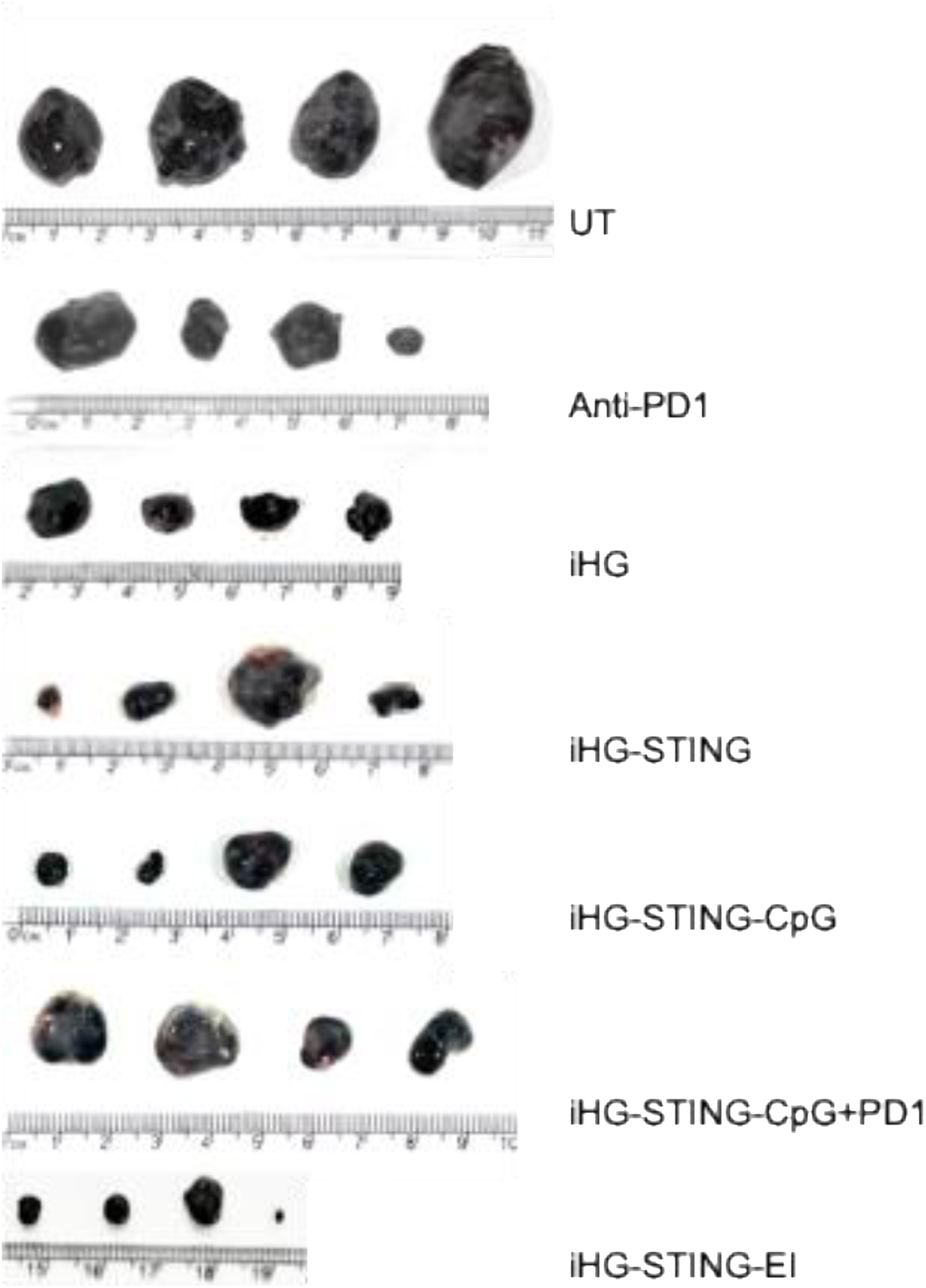
Tumor images after 21 days post-B16F10 tumor induction for different treatment groups.

