## supplementary figures for "Injectable Immune-Engineered Hydrogel Niche Remote From The Immune Suppressed Tumor Microenvironment For Cancer Immunotherapy"

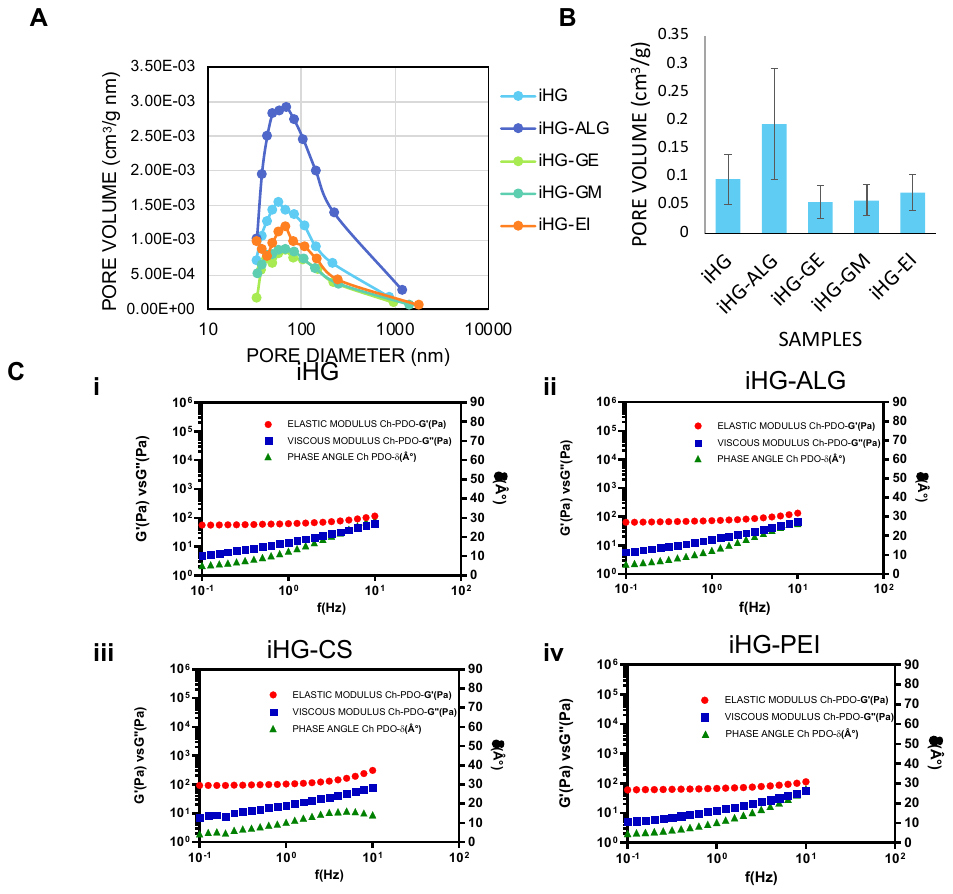

Figure S1 **Characterisation of Immunohydrogel (iHG)** (A) BJH pore size distribution curves for iHGs with varying compositions, highlighting differences in pore size and volume across samples. (B) Bar graph showing the average pore volume of approximately 0.1 cm^3^/g for iHGs with different compositions, derived from BET analysis for iHGs with different compositions. (C) Rheological analysis of gels with varying compositions (i–iv). Frequency sweep curves demonstrate the shear-thinning behavior and consistent flow properties of the gels.

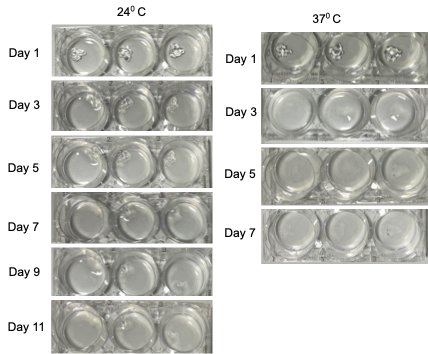

Figure S2**. In vitro degradation study of iHG gel.** (A) Representative images showing gradual degradation of the gel over 11 days in 1× PBS at 24 °C, indicating slow dissolution. (B) Faster degradation observed within 7 days in 1× PBS at 37 °C.

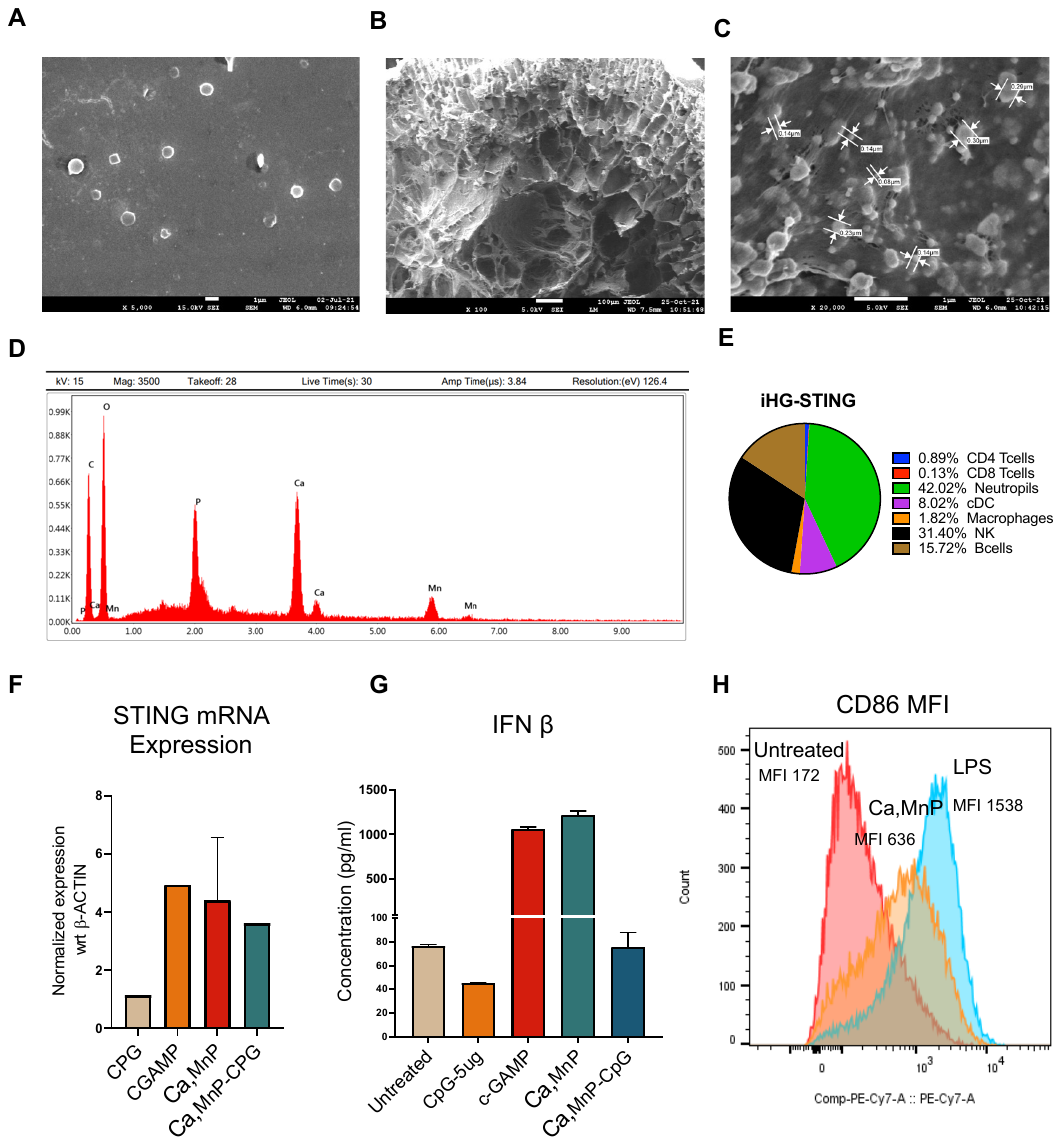

Figure S3**. iHG STING characterization.** (A) SEM images of (Ca, Mn)P nanoparticles showing multidispersered nanoparticles with size range from 1000μm to 100nM. (B) SEM images (100x) of a lyophilized section of iHG loaded with STING activating nanoparticles. (C) High magnified images of (Ca, Mn)P nanoparticle loaded within iHG gel. (D) EDAX spectrum of loaded nanoparticle showing a peak of calcium and manganese within iHG. (E) Immune cell infiltration in iHG-STING gel post 5 days after subcutaneous implantation in C57BL/6J mouse, showing a high infiltration of neutrophils, NK cells, B cells, and DC. (F)shows the increase in the mRNA expression level of STING after 24 hrs treatment 100μM (Ca, Mn)P and (Ca, Mn)P+CpG in the JAWS II DC cell line. (G) Shows the release of type I interferon (IFN-$\beta$) by STING activation by JAWS II DC cell line with the treatment of 100μM (Ca, Mn)P and (Ca, Mn)P+CpG. (H) Increase in CD86 activation (MFI) in JAWS II DC with treatment with 100μM (Ca, Mn)P nanoparticle for 24 hours.

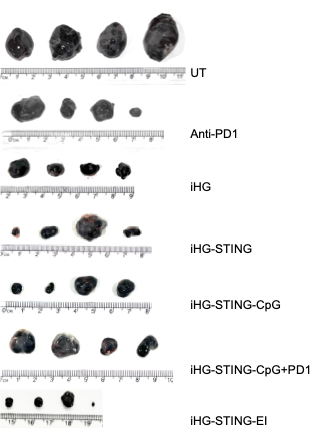

Figure S4**. Tumor images** after 21 days post-B16F10 tumor induction for different treatment groups.
